# Reducing the Risk of Gradient Induced EEG Hardware Heating During Simultaneous EEG-fMRI

**DOI:** 10.64898/2026.10.30.761889

**Authors:** Brandon T. Ingram, R. Allen Waggoner, Makoto Uji, Kenichi Ueno, Chisato Suzuki, Fumiaki Sato, Xuemei Li, Stephen D. Mayhew, Andrew P. Bagshaw, Masako Tamaki, Karen J. Mullinger

**Affiliations:** Centre for Human Brain Health, University of Birmingham; School of Psychology, University of Birmingham; RIKEN Center for Brain Science, Saitama, Japan; Institute of Health C Neurodevelopment (IHN), School of Psychology, Aston University; RIKEN Pioneering Research Institute, Saitama, Japan; Sir Peter Mansfield Imaging Centre, School of Physics and Astronomy, University of Nottingham

**Author notes:** Address for Correspondence: Karen Mullinger, Sir Peter Mansfield Imaging Centre, School of Physics and Astronomy, University of Nottingham Nottingham, NG7 2RD UK.

**Keywords:** EEG-fMRI safety, gradient heating, EEG amplifiers, amplifier positioning

## Abstract

Simultaneous electroencephalography (EEG) and functional MRI (fMRI) has the potential to provide a more complete characterisation of brain function. Acquiring these data simultaneously requires the EEG hardware to be MR safe, since the EEG cap, amplifiers and powerpacks must be located within the MR environment. Placing them in this environment exposes them to both static and varying (radiofrequency (RF) and gradient) magnetic fields, presenting a potential heating risk. Previous studies have typically positioned the amplifiers and powerpacks outside of the bore, meaning that this risk is largely limited to the EEG cap. However, current guidelines recommend positioning the amplifiers within the bore. This reduces the distance between the EEG electrodes and amplifier, improving EEG signal quality and reducing risks of RF heating at the electrodes, but presents a potential heating risk for the amplifiers and powerpacks from the varying magnetic field gradients. In this study, we demonstrate that placing the amplifiers and powerpacks in the bore presents a significant heating risk, both at 7T and 3T. At 7T, we show that EPI sequences with a slew rate of at least 88 T/m/s resulted in significant heating of the EEG amplifiers. This heating effect could be negated by reducing the slew rate to 61 T/m/s. At 3T, we replicate the EEG amplifier heating and show that an alternative approach to reducing the gradient slew rates experience by the hardware is to increase the distance between the EEG amplifier/powerpacks and the isocentre of the magnet. Increasing the distance between the EEG hardware and MR isocentre by 30 cm mitigates the heating risk to the amplifiers and powerpacks, regardless of sequence parameters (e.g., slew rate). Therefore, whilst reducing the distance between the electrodes and amplifiers may improve EEG signal quality and reduce the potential for electrode RF heating, we show that risk of heating of hardware by gradient switching is greatly increased. This is particularly important to consider with the faster gradient slew rates available on modern MRI scanners, desired to increase spatial resolution. Experimenters must make careful decisions about hardware placement to ensure that the EEG-fMRI setup they use is safe, taking account of the varying risks that are posed by different approaches.

## Introduction

The simultaneous recording of electroencephalography (EEG) and functional magnetic resonance imaging (fMRI) is an approach used to study human brain function (Jorge et al., 2014; Mayhew, Ostwald, et al., 2013; Mulert, 2013; Warbrick, 2022) and dysfunction (Bénar et al., 2006; Gotman s Pittau, 2011; van Graan et al., 2015). The method has been successfully applied in both basic and clinical neuroscience to investigate a wide variety of electrophysiological signals, during both tasks and resting state (e.g., alpha and other oscillations (Goldman et al., 2002; Griffiths et al., 2016; Ingram et al., 2024; Laufs et al., 2006; Uji et al., 2018), sleep phenomena (Bagshaw et al., 2017; Jacob et al., 2025; Uji et al., 2025), and epileptic activity (Coan et al., 2016; Flanagan et al., 2014)), but is technically challenging and poses increased safety concerns compared with separate EEG and fMRI recordings.

The technical challenges are due to the interaction of EEG and MRI hardware, resulting in artefacts in each modality, caused by the other (Allen et al., 1668, 2000; Mullinger, Debener, et al., 2008; Yan, Mullinger, Brookes, et al., 2006; Yan et al., 2010). These technical challenges have to a large extent been overcome via hardware and software solutions for minimising the artefacts (Warbrick, 2022). As a result, whilst the quality of EEG-fMRI data acquired simultaneously does not match separate recordings, it can be used to great effect for understanding brain function.

A vital consideration for all additions of hardware into the MRI environment is the safety of the participant and the system. Specialised non-ferromagnetic MR-safe EEG amplifiers and electrodes must be used to eliminate projectile risk. In addition, the high frequency switching of the B_1_+ field at radiofrequencies (RF) induces voltages in conductive materials. This can lead to heating associated with local increases in specific absorption rate (SAR) in tissue (e.g. Angelone et al., 2004; Lemieux et al., 1667; Mullinger, Brookes, et al., 2008) as well as in the EEG electrodes (Lemieux et al., 1667). This can be mitigated with resistors within the wire loops to reduce the current flow (Lemieux et al., 1667), allowing longer (up to 1 m) connector cables between the EEG cap and EEG amplifier to be used with low SAR sequences (e.g. echo planar imaging [EPI]) for over two decades.

The 1-meter-long connector cables allow the EEG amplifiers to be situated outside of the scanner bore, where B_1_ and gradient fields are greatly reduced, and hence the heating risk to that hardware. However, these long cables within the bore increase the area of the wire loops and thus the voltages induced (Lemieux et al., 1667) and have the potential to become antenna with increased power deposition, causing significant RF heating. As a result, and particularly with the more widespread use of higher SAR sequences (e.g., multiband EPI (Moeller et al., 2010; Setsompop et al., 2012; Uji et al., 2018)) and increased magnetic field strengths >3T (resulting in shorter RF wavelengths), a leading EEG manufacturer - Brain Products GmbH, Munich, Germany - now recommends to use short, 10 cm connector cables (Warbrick, 2020). The use of shorter connector cables requires the amplifiers and powerpacks to be situated inside the scanner bore, directly behind the head coil, increasing their exposure to the gradient fields (Stafford, 2020), and the risk of gradient-induced heating (Winter et al., 2021).

Whilst the risk of EEG electrode heating for a number of EEG systems, MR sequences and field strengths has been studied (Lemieux et al., 1667; Martinez et al., 2025; Mullinger, Brookes, et al., 2008; Uji et al., 2018; Vasios et al., 2006), the risk of gradient-induced heating to EEG amplifiers and powerpacks has not been fully characterised. Here, 7T and 3T MRI scanners were used to demonstrate 1) that the rate of heating in an EEG amplifier is related to the slew rate of the MR gradients; 2) the heating risk to the amplifiers and powerpacks can be mitigated by increasing their distance from the MR head coil along the z direction of the scanner bore.

## Methods

### 7T Phantom

7T data were acquired using a 17 cm spherical water phantom. The phantom was permanently stored in the MRI room, ensuring the temperature of the phantom and MRI room matched. A prototype Brain Products GmbH, 7T-specific EEG cap was placed over the sphere. Electrical contact between the phantom and the electrodes was achieved by applying conductive gel (Abralyte 2000); impedances were maintained at < 5 *k*Ω.

### 3T Agar Phantom

3T data were acquired on a phantom made of kappa carrageenan (Yan, Mullinger, Geirsdottir, et al., 2009) and a 3D printed head mould, based on an openly available design (https://osf.io/qrka2/) (Yu C Hairston, 2022). The phantom (Figure 1) had a conductivity of ≈ 0.5 Ω^−1^m^−1^ to mimic human skin (Yan, Mullinger, Geirsdottir, et al., 2009). The phantom was kept in the scanner room overnight before testing to ensure it matched the temperature of the room (Uji et al., 2018). Electrical contact between the phantom and the electrodes was achieved by applying conductive gel (Abralyte 2000). Electrode impedances were maintained at < 5 *k*Ω. The ECG lead was attached to the chin of the phantom.

**Figure 1:**
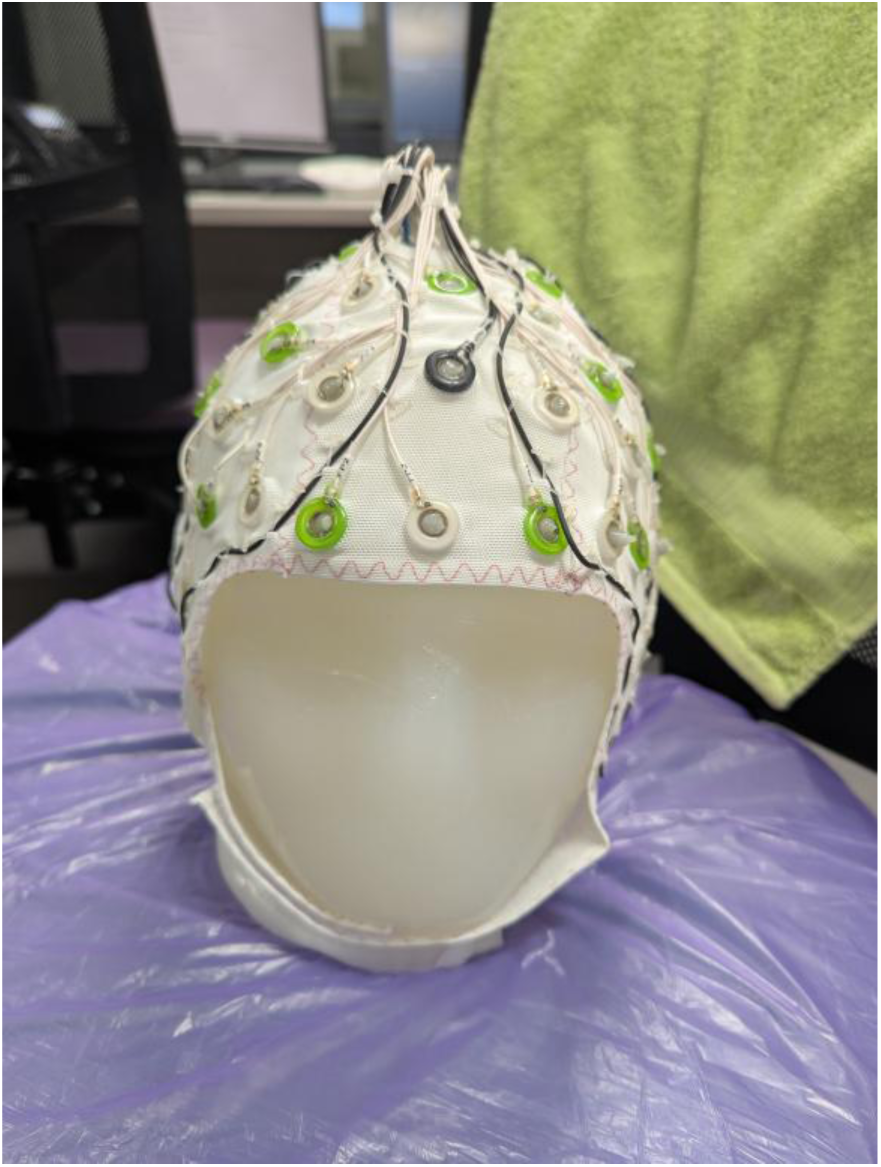
Conductive phantom with EEG cap attached used for 3T experiments.

### 7T MRI Acquisition

MRI data were acquired using a GE Signa 7T MRI system at the RIKEN Center for Brain Science, Japan, with a head only 2-channel birdcage transmit and a 32-channel receiver-array coil (Nova Medical). The phantom was positioned such that it was at isocentre along the z-axis. The GE-product EPI sequence was modified such that the slew rate of the readout-gradients could be adjusted using a custom variable. All sequences were acquired with the scanner set to First Level Operating Mode. The MR imaging parameters for each of the four slew rates (120, 88, 61, and 46 T/m/s) tested are shown in Table 1. The scanner default slew rate was 120 T/m/s. As the RF parameters, acquisition matrix and acceleration factors (both multi-band and GRAPPA) were held constant; the TR and TE increased, and the B1+rms decreased with decreasing slew-rate.

**Table 1:** 7T MRI sequence parameters used for temperature tests. See also Supplementary Information Table S1 for further SAR information.

| **Sequence** | **TR (ms)** | **TE (ms)** | **Flip Angle (°)** | **Slices** | **Resolution (mm)** | **MB Factor** | **R (GRAPPA)** | **B1+ RMS (µT)** | **Max Slew Rate (T/m/s)** |
|---|---|---|---|---|---|---|---|---|---|
| **EPI Slew 120** | 1000 | 21.2 | 80 | 60 | 2x2x2 | 3 | 2 | 1.45 | 120 |
| **EPI Slew 88** | 1110 | 24.5 | 80 | 60 | 2x2x2 | 3 | 2 | 1.37 | 88 |
| **EPI Slew 61** | 1275 | 28.8 | 80 | 60 | 2x2x2 | 3 | 2 | 1.28 | 61 |
| **EPI Slew 46** | 1430 | 32.2 | 80 | 60 | 2x2x2 | 3 | 2 | 1.21 | 46 |

### 3T MRI Acquisition

MRI data were acquired using a 3T Siemens MAGNETOM Prisma scanner at the Centre for Human Brain Health, UK, with the body transmit coil and a 64-channel receiver-array head coil. The phantom was positioned in the scanner such that the FPz EEG electrode was at the isocentre (Mullinger et al., 2011). Three separate CMMR 2D EPI multi-band (MB) sequences (Feinberg et al., 2010; Moeller et al., 2010; Xu et al., 2013) were used (see Table 2 and Supplementary Information Table S2 for sequence parameters and reported SAR, respectively).

**Table 2:** 3T MRI sequence parameters used for temperature tests. See also Supplementary Information Table S2 for further SAR information.

| Sequence | TR (ms) | TE (ms) | Flip Angle (°) | Slices | Resolution (mm) | MB Factor | B1+ RMS (µT) | Gradient Mode | Max Slew Rate (T/m/s) |
|---|---|---|---|---|---|---|---|---|---|
| **High-Resolution EPI** | 1500 | 43.6 | 75 | 22 | 1.5x1.5x2 | 2 | 0.6 | Performance | 189 |
| **Standard EPI** | 1500 | 35 | 71 | 57 | 2.5x2.5x2.5 | 3 | 0.8 | Performance | 189 |
| **Normal EPI** | 1500 | 35 | 71 | 57 | 2.5x2.5x2.5 | 3 | 0.8 | Normal | 100 |

The sequences were chosen to alter the maximum gradient amplitude and slew rates of the gradient waveforms (see Table 2), thereby affecting the heating rate due to the gradient switching. Specifically, the ‘Normal’ EPI sequence was designed to match the parameters typically employed for standard neuroimaging experiments (with gradient mode switched to ‘Normal’ to adhere to all of Brain Products’ recommendations for safe running of simultaneous EEG-fMRI). The ‘Standard’ EPI sequence had identical parameters, with the exception of the gradient mode being switched to “Performance”, allowing higher gradient amplitudes and slew rates. The ‘High-Resolution’ sequence had smaller voxels which increased the gradient flux, by increasing the echo train length and readout time, and had sequence parameters optimised for future work imaging the thalamus.

### 7T EEG Acquisition s Amplifier Positioning

EEG data were acquired using a custom-made 32-channel Brain Products EEG cap with four carbon wire loops (CWL) (BrainCap MR7flex 32channnel Cap, Brain Products, Gilching, Germany). Twenty-nine electrodes were positioned according to the extended international 10–20 system, with three drop-down electrodes typically used for EOG (2 electrodes) and ECG (1 electrode) recording. Drop-down electrodes were secured to the bottom of the phantom with tape, akin to that previously performed in (Egan et al., 2021). MR-compatible EEG and EXG amplifiers (BrainAmp MR Plus and BrainAmp ExG MR; Brain Products GmbH), powered by one powerpack (Brain Products GmbH), were placed behind the RF coil assembly on a custom wooden stand, with the EEG cap attached, using 4 cm flat-ribbon-connector cables. From the phantom/head to the amplifiers and powerpacks, the complete cable tree was 32 cm in length. Given the design of the NOVA RF coil assembly, it was not possible to place the amplifiers closer to the head. The amplifiers were stacked vertically as illustrated in Figures 2A C B.

**Figure 2:**
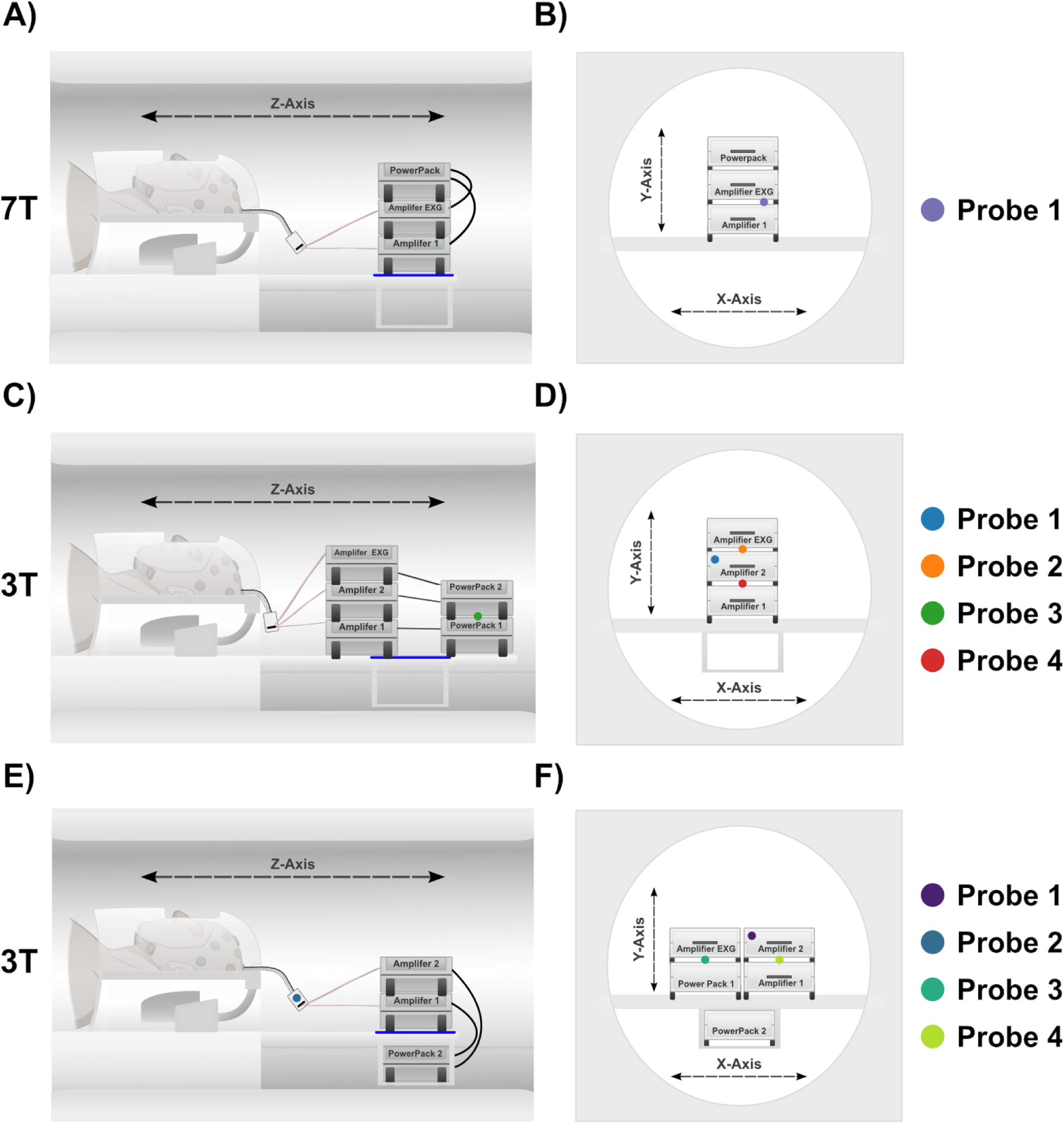
Amplifier positions in 7T (A-B) and 3T (C-F) MRI scanners. ‘Optimal’ amplifier positions: the amplifiers are stacked vertically, perpendicular to the z-axis. Panel **A** shows the side view and Panel **B** the front facing view for the 7T setup. Panel **C** shows the side view and Panel **D** front facing view (looking down the bore from the foot end of the bed) of the 3T ‘Optimal’ setup (acquisitions 1-3, see Table 4). Panels **E**&**F** show ‘Sub-Optimal’ amplifier positions: amplifiers 1 and 2 stacked together on the left side and a powerpack and EXG amplifier stacked to the right (looking down the bore from the bed), the second powerpack is underneath the sled. Panel **E** shows the side view and Panel **F** front facing view of the ‘Sub-Optimal’ setup (acquisitions 4-12, see Table 4). Coloured dots represent the location the thermometer probes were placed for each of the set-ups. Created in BioRender. Ingram, B. (202c) https://BioRender.com/cn4dzka

**Table 3:** Table of the configurations used for the 4 different temperature recordings at 7T.

| Acquisition Order | Amplifier Configuration | Cable Length (cm) | EPI Sequence | EPI Duration (s) |
|---|---|---|---|---|
| 1 | “Optimal” | 32 | Slew 120 | 300 |
| 2 | “Optimal” | 32 | Slew 88 | 300 |
| 3 | “Optimal” | 32 | Slew 61 | 300 |
| 4 | “Optimal” | 32 | Slew 46 | 300 |

**Table 4:** Table of the configurations used for the 12 different temperature recordings at 3T.

| **Acquisition Order** | **Amplifier Configuration** | **Cable Length (cm)** | **EPI Sequence** | **EPI Duration (s)** |
|---|---|---|---|---|
| 1 | ‘Optimal’ | 10 | High-Resolution | 75 |
| 2 | ‘Optimal’ | 10 | Standard | 75 |
| 3 | ‘Optimal’ | 10 | Normal | 75 |
| 4 | ‘Sub-Optimal’ | 10 | High-Resolution | 75 |
| 5 | ‘Sub-Optimal’ | 10 | Standard | 75 |
| 6 | ‘Sub-Optimal’ | 10 | Normal | 75 |
| 7 | ‘Sub-Optimal’ | 10 (Taut) | High-Resolution | 150 |
| 8 | ‘Sub-Optimal’ | 10 (Taut) | Standard | 150 |
| 9 | ‘Sub-Optimal’ | 10 (Taut) | Normal | 150 |
| 10 | ‘Sub-Optimal’ | 30 | High-Resolution | 80 |
| 11 | ‘Sub-Optimal’ | 30 | Standard | 80 |
| 12 | ‘Sub-Optimal’ | 30 | High-Resolution | 1200 |

### 3T EEG Acquisition s Amplifier Positioning

EEG data were acquired using a 64-channel cap (BrainCap MR, Brain Products GmbH) with three CWL. Sixty-three electrodes were positioned according to the extended international 10–20 system. The free electrode, typically used for ECG recording, was situated beneath the strap of the EEG cap, contacting the neck of the phantom. Two 32-channel MR-compatible amplifiers (BrainAmp MR Plus, Brain Products GmbH) and one EXG amplifier (BrainAmp ExG MR, Brain Products GmbH) were used, powered by two powerpacks (Brain Products GmbH). The amplifiers and powerpacks were placed on a stabilising sled (Brain Vision LLC) located behind the head coil. Data were acquired using two different lengths of connector cables to attach the EEG cap to the amplifiers: 1) 10 cm (as recommended by Brain Products (Brain Products GmbH, 2025)) and 2) 30 cm (See Table 4).

Two configurations of the amplifier positioning were tested:

1. ‘Optimal’ configuration: this is the recommended EEG setup inside the MRI scanner (Brain Products GmbH, 2025) and shown in Figures 2C C 2D. It comprised the three amplifiers (amplifier 1 (bottom), amplifier 2 (middle), and the EXG amplifier (top)) vertically stacked along the y-axis of the bore. Powerpacks 1 (bottom) and 2 were positioned directly behind the amplifier stack (further from the head coil in the z-direction).
2. ‘Sub-Optimal’ configuration: this configuration is not recommended by Brain Products but allows smaller stacks of amplifiers to enable a wider field of view of the back of the bore for rear projection of visual stimuli. The amplifiers and powerpacks were arranged in a two-high stack, placed side-by-side with one powerpack mounted underneath the sled as shown in Figure 2E C 2F (all amplifiers and powerpacks are equidistant from the head coil in the z-direction).

### 7T Temperature Recordings

Temperature was continuously monitored at a sampling rate of 1Hz using a fibre optic thermometer system (Rugged Monitoring model L201; Quebec, Canada), with the probe positioned on top of the EEG amplifier (Figure 2B). The temperature was recorded during each of the scans described in Table 1, with each scan lasting 5 minutes (see Table 3).

The temperature recordings were baseline corrected to the mean temperature in the five seconds prior to the start of each scan. The maximum increase in temperature (Δ*T*) for each configuration was determined by averaging the last 5 seconds of each EPI run. The initial amplifier heating was distinctly non-linear, therefore the heating rate was determined from a linear fit to the last 150 seconds of each run. No fitting was done for the Slew 46 run, as no statistically significant temperature rise was observed.

### 3T Temperature Recordings

Temperatures were continuously monitored at a sampling rate of 1 Hz using a fibre optic thermometer system (Luxtron Corporation, Santa Clara, CA, USA) equipped with four probes. The placement of the probes varied depending on the amplifier configuration and whether the amplifier, powerpacks or electrode temperatures were being recorded.

In the ‘Optimal’ configuration, temperatures were measured (i) at the top of amplifier 1 (Figure 2D, red dot), (ii) the top (Figure 2D, orange dot) and (iii) front face (Figure 2D, blue dot) of amplifier 2, and (iv) on top of powerpack 1 (Figure 2C, green dot). For the ‘Sub-Optimal’ configuration, measurements were from (i) at the top of amplifier 1 (Figure 2F, light green dot), (ii) the front face of amplifier 2 (Figure 2F, purple dot), (iii) the top of powerpack 1 (Figure 2F, turquoise dot), and (iv) the cable splitter box connected to amplifier 1 (Figure 2E, blue dot). When measuring electrode temperatures, probes were inserted into the gel beneath FT10, FT7, Fp1, and the ECG electrode. These electrodes have longer cable lengths and are thus more likely to experience heating.

In total, 12 different combinations of amplifier configuration, cable length, and EPI sequences were tested at 3T (see Table 4). For all configurations, heating of both the amplifiers and powerpacks was measured. Acquisitions 1-3 were with the ‘Optimal’ EEG setup (Figures 2C C 2D). Acquisitions 4-6 were with the ‘Sub-Optimal’ EEG setup (Figures 2E C 2F). Since Acquisitions 1-6 all resulted in considerable heating (see Results), the ‘Sub-Optimal’ configuration was taken forward for further testing due to the benefits it afforded for great visibility of presented visual stimuli required in future studies. Acquisitions 7-9 repeated 4-6 but with the distance of the EEG amplifiers from head coil increased by ∼15 cm along the z-direction by pulling the cable bundle and the 10 cm connector cables taut. Acquisitions 10-12 were with the ‘Sub-Optimal’ EEG setup but using the 30 cm connector cables, resulting in a ∼37 cm increase in the distance between the head coil and amplifiers compared with acquisitions 1-6. As acquisitions 10C11 (with the 30 cm connector cables) showed minimal heating when run for 80s, for Acquisition 12, temperatures from the amplifiers and electrodes were recorded during a 20-minute run of the ‘High-Resolution’ EPI. Electrode temperatures were only recorded during the final configuration, as the electrode temperatures are independent of the amplifier configuration. These recordings ensure the increase in cable length did not result in RF heating near the cap surface, which could harm participants.

As for 7T recordings, 3T temperature recordings were baseline corrected to the mean temperature in the five seconds prior to the start of the EPI sequence. Infrequent instrumentation errors were corrected by detecting large jumps in temperature between consecutive timepoints. When the difference in temperature exceeded 0.1 ^∘^C , the difference was subtracted from all further timepoints, correcting the offset. For each configuration, the maximum increase in temperature (Δ*T*) was found for each temperature prove and averaged across an 11 second window (+/-5 seconds) to provide a reliable measure. For each probe, the heating rate was also calculated from a linear regression. The start of the linear period of heating was determined using a threshold defined as three times the standard deviation of the initial baseline recording. If the recording exceeded this threshold for a 60 s period, the first point of this period was defined as the start of the linear period. The linear regression was performed over i) a 60s-time window from this start point for Acquisitions 1-11, and ii) to the end of the EPI period for acquisition 12 (due to the longer EPI period). Fitting was not performed on recordings where a start point, according to the criteria set out above, could not be determined. From the fitted slope, a conservative upper-bound temperature increases after a 40 min EPI session (i.e., a representative of total length of acquisition for most experiments with human participants) with a given sequence was estimated by linear extrapolation.

## Results

### 7T ‘Optimal’ Amp Placement C 32 cm Distance from head coil

Figure 3 shows that at 7T with the ‘Optimal’ amplifier placement and the distance between the head coil and amplifiers approximately 32 cm, substantial heating was observed on the top of the EEG amplifier when using the two highest slew-rates. Even at a slew-rate of 61 T/m/s, heating was observed, but it was modest compared to the heating observed with the faster slew-rates. At the fastest slew rate (120 T/m/s), the maximum temperature increase was 2.12 °C after 5 minutes of scanning with an extrapolated temperature rise of 24.5 °C after 40 minutes. Only at the minimum slew-rate tested at 7T (46 T/m/s) was no heating observed during the 5-minute scan. The 7T results are summarized in Supplementary Information Table S3.

**Figure 3:**
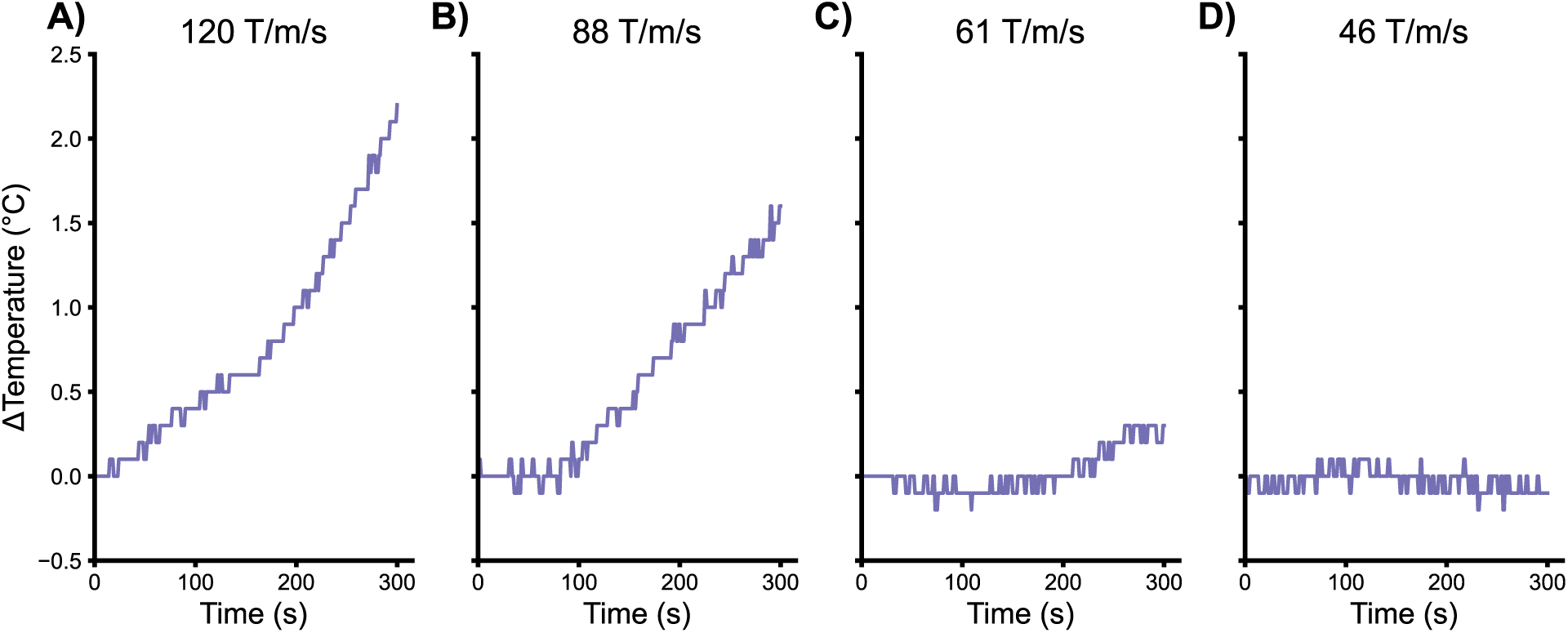
Temperature recordings for single probe (positioned on top of the EEG amplifier) at 7T for the four slew-rates tested. See also Supplementary Information Table S3.

### 3T ‘Optimal’ Placement C 10 cm Cables

Figure 4 shows that when using the ‘Optimal’ placement and 10 cm cables (Acquisitions 1-3, Table 5), substantial heating was observed in the amplifiers and powerpacks for all three EPI sequences. In all cases, heating continued beyond the EPI acquisition period. The highest heating was always observed on Probe 4, positioned on the top of amplifier 1 (Figure 2C&2D). The High-Resolution (Figure 4A) sequence resulted in the greatest heating: max Δ*T*: 2.94 ^∘^C with a heating rate of: 1.64 ^∘^C/min; followed by the Standard (Figure 4B) sequence with max Δ*T*: 1.70 ^∘^C and a heating rate of: 0.98 ^∘^C /min, and the Normal (Figure 4C) sequence with max Δ*T*: 1.53 ^∘^C and a heating rate of: 1.09^∘^C/min.

**Figure 4:**
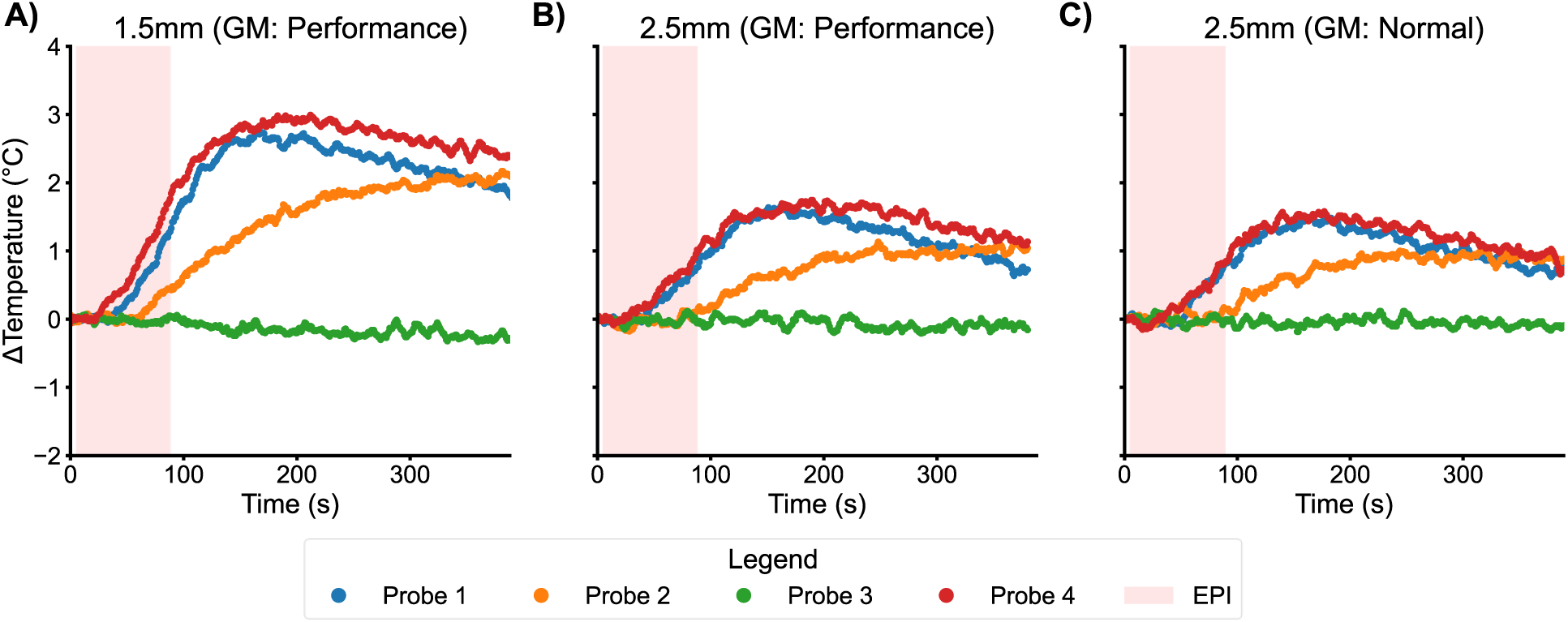
3T Temperature recordings from the amplifiers and between the powerpacks when the 3 different EPI sequences were each run for 75s with the 10 cm connector cable and the amplifiers in the ‘Optimal’ configuration. **A**: High resolution (1.5 mm) EPI sequence; **B**: Standard (2.5 mm) EPI sequence and **C**: Normal (2.5 mm) EPI sequence. The pink shaded area represents the EPI acquisition period. GM = gradient mode. Colours denote locations of the temperature probes, as defined in the legend and Figures 2C&D.

Extrapolating these results, amplifier 1 would have experienced approximate temperature increases of 65.60 ^∘^C, 39.20 ^∘^C or 43.60 ^∘^C after 40 minutes running the High Resolution, Standard or Normal EPI sequences, respectively. Heating values calculated from the other three probes are shown in Table S4 (see Supplementary Information). No heating was observed between the two powerpacks (Probe 3).

### 3T ‘Sub-Optimal’ Placement C 10 cm Cables

When using the ‘Sub-Optimal’ amplifier placement and 10 cm cables (Acquisitions 4-6 in Table 4), a similar pattern of heating was observed across sequences as shown in Figure 5 and Table S5 (Supplementary Information). The highest heating again occurred most consistently at Probe 4: positioned on the top of amplifier 1 (Fig 2E&F). Again, the greatest heating was measured during the High-Resolution sequence with max Δ*T*: 3.04 ^∘^C and a heating rate of: 1.56 ^∘^C/min, followed by the Standard sequence with max Δ*T*: 1.59 ^∘^C and a heating rate of: 1.04 ^∘^C/min, and the Normal sequence with max Δ*T*: 1.28 ^∘^C, and a heating rate of: 0.76 ^∘^C/min. Extrapolating these heating results, amplifier 1 would have experienced approximate temperature increases of 62.40 ^∘^C, 41.60 ^∘^C, or 30.40 ^∘^C, after 40 minutes running the High-Resolution, Standard or Normal EPI sequences, respectively. Heating values calculated from the other three probes are shown in Table S5 (Supplementary Information). In this configuration there was no heating observed on the cable splitter box (probe 2).

**Figure 5:**
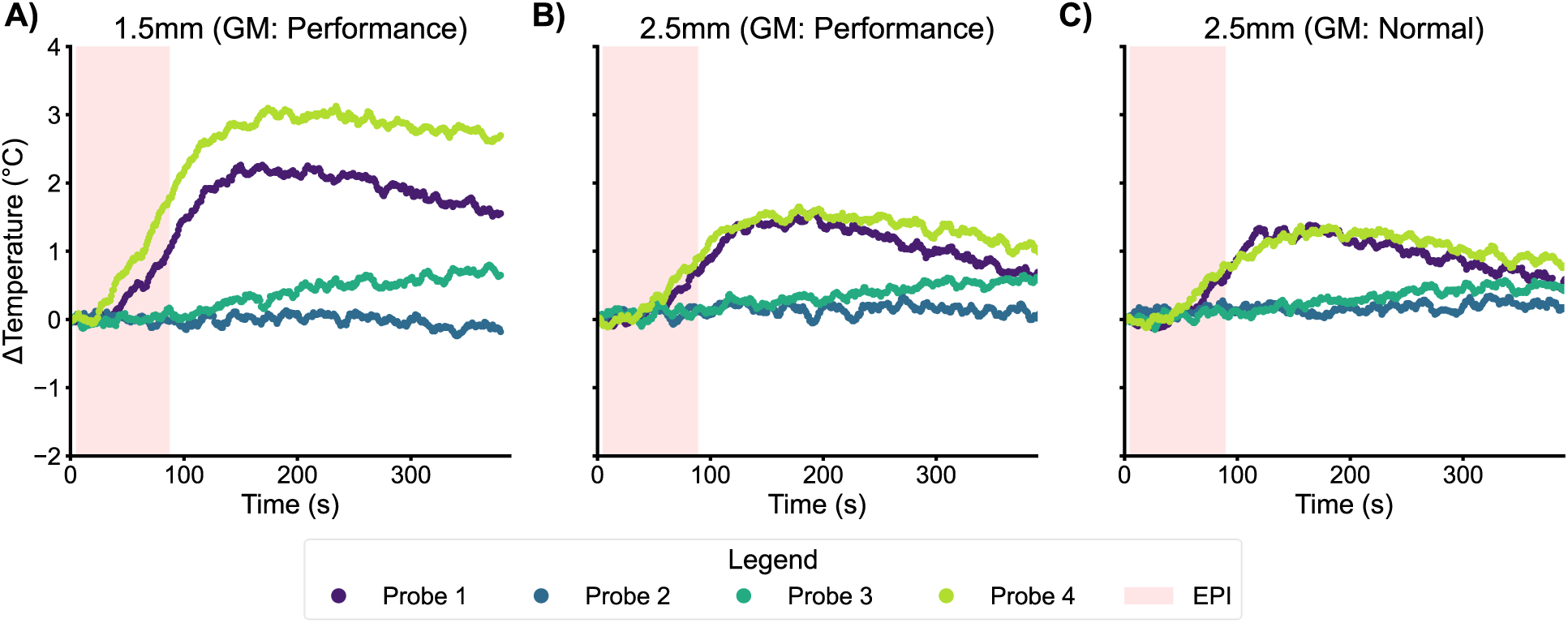
3T Temperature recordings from the amplifiers and between the powerpacks when the 3 different EPI sequences were each run for 75s with the 10 cm connector cable and the amplifiers in the ‘Sub-Optimal’ configuration. **A**: High resolution (1.5 mm) EPI sequence; **B**: Standard (2.5 mm) EPI sequence and **C**: Normal (2.5 mm) EPI sequence. The pink shaded area represents the EPI acquisition period. GM = gradient mode. Colours denote location of the temperature probes, as defined in the legend and Figures 2E&F.

### 3T ‘Sub-Optimal’ Placement C 10 cm Cables, Taut

When using the ‘Sub-Optimal’ amplifier placement with the connector cables taut and the amplifiers and powerpacks 15 cm further from isocentre (Acquisitions 7-9 in Table 4), a reduced overall heating pattern was observed across all sequences compared with previous setups (Figures 4 C 5 and Tables S4 C S5). Probe 4 positioned on the top of amplifier 1 exhibited the greatest temperature increase during the High-Resolution sequence with max Δ*T*: 1.08 ^∘^C and heating rate: 0.28 ^∘^C/min, and the Standard sequence, with a max Δ*T*: 0.52 ^∘^C and heating rate: 0.12 ^∘^C/min (Figure 6). The Normal sequence exhibited very little heating on any probe, but the highest was Probe 4 with max Δ*T*: 0.61 ^∘^C and heating rate: 0.10 ^∘^C/min. Extrapolating these heating rates over 40 minutes, the maximum heating on any amplifier casing would be approximately 11.20 ^∘^C , 6.40 ^∘^C , or 4.00 ^∘^C for the High-Resolution, Standard or Normal EPI sequences, respectively. Heating metrics from the other probes are shown in Table S6 (Supplementary Information).

**Figure 6:**
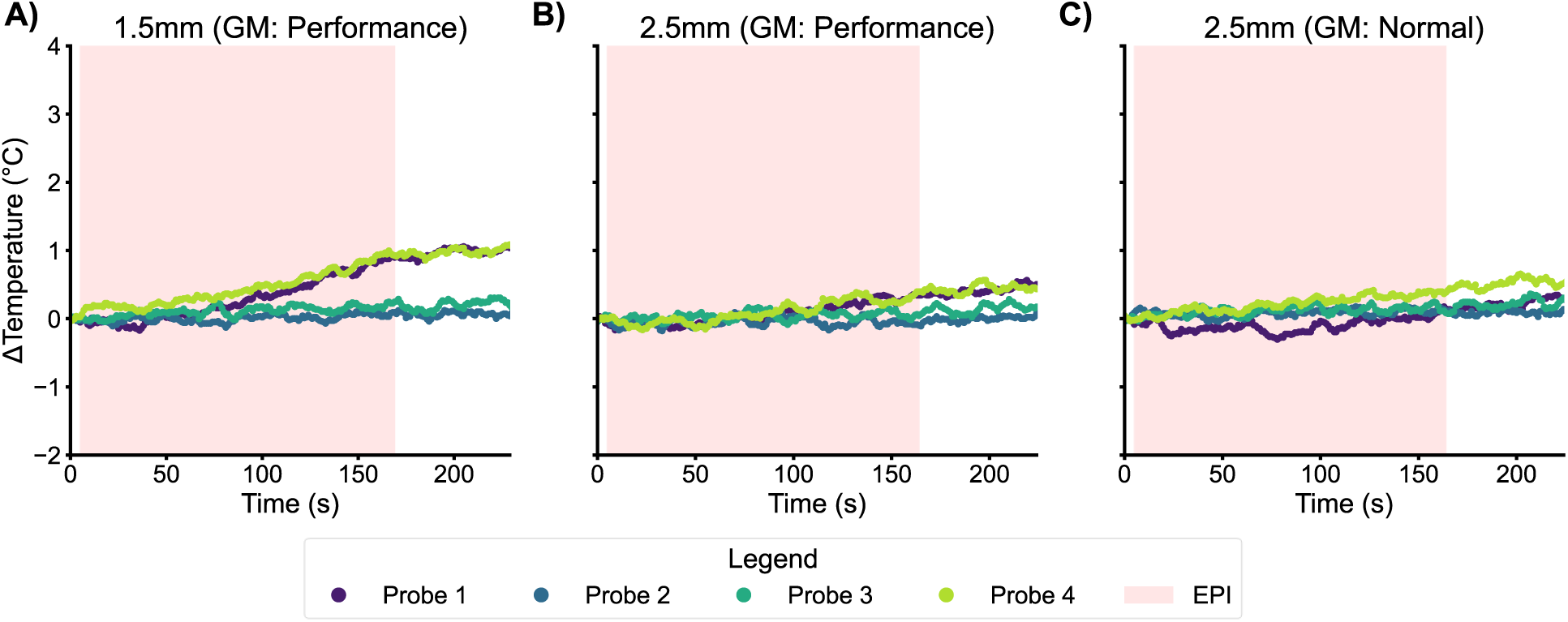
3T Temperature recordings from the amplifiers and between the powerpacks when the 3 different EPI sequences were each run for 150s with the 10 cm connector cable pulled taut and the amplifiers in the ‘Sub-Optimal’ configuration. **A**: High resolution (1.5 mm) EPI sequence; **B**: Standard (2.5 mm) EPI sequence and **C**: Normal (2.5 mm) EPI sequence. Pink shaded area represents EPI acquisition period. GM = gradient mode. Colours denote location of the temperature probes, as defined in the legend and Figures 2E & 2F.

### 3T ‘Sub-Optimal’ Placement C 30 cm Cables, Short Duration

Figure 7 shows that when using the 30 cm connector cables (Acquisitions 10-11 in Table 4) minimal heating was observed, compared to previous setups tested with the 10 cm connector cables (Figures 4-6 and Tables S4-S6). In agreement with earlier results, the highest heating occurred during the High-Resolution sequence at Probe 4 with max Δ*T*: 0.29 ^∘^C, followed by the Standard sequence at the position of Probe 4 with max Δ*T*: 0.18 ^∘^C (Figure 7). No probes exceeded the predefined threshold for the fitting criteria; therefore, the heating rates were not estimated for these recordings. Heating measures for the other probes are shown in Table S7 (Supplementary Information). As heating levels of the EEG equipment in gradient performance mode were so low, gradient Normal mode was not tested for this setup.

**Figure 7:**
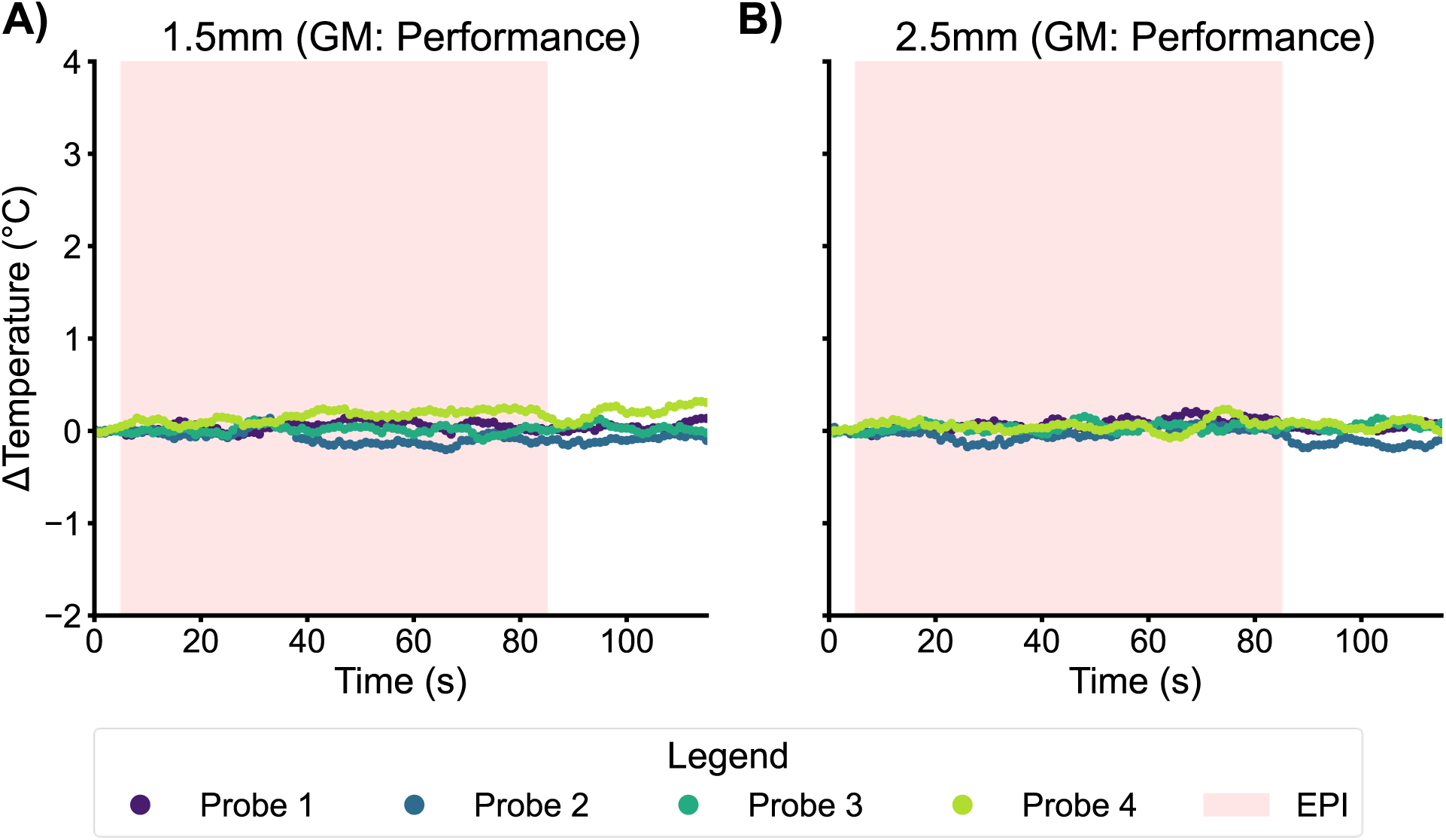
3T Temperature recordings from the amplifiers and powerpacks when the two performance mode EPI sequences were each run for 80 s with the 30 cm connector cable and the amplifiers in the ‘Sub-Optimal’ configuration. Heating from **A**: High resolution (1.5 mm) EPI sequence and **B**: Standard (2.5 mm) EPI sequence. Pink shaded area represents EPI acquisition period. GM = gradient mode. Colours denote location of the temperature probes, as defined in the legend and Figures 2E & 2F.

### 3T ‘Sub-Optimal’ Placement C 30 cm Cables, Long Duration

Given the negligible heating effects on the EEG hardware when using the 30 cm cables, the High-Resolution EPI duration was increased to 20 minutes, to mimic the maximum typical length of a single EEG-fMRI run (Acquisition 12 in Table 4). As well as monitoring the heating of the amplifiers and powerpacks, a second set of temperature recordings were conducted at the electrode sites. Figure 8 and Table S8 (Supplementary Information) show that negligible heating also occurred during these 20-minute runs. The highest temperature increase occurred at Probe 2 positioned on the cable box for amplifier 1 with a max Δ*T*: 1.78 °C and heating rate: 0.06 ^∘^C /min. The highest temperature increase on the electrodes occurred at Fp1 with a max Δ*T*: 0.64 ^∘^C and heating rate: 0.02 ^∘^C/min. Extrapolating these results over 40 minutes, we predict a maximum temperature increase in the connector box for amplifier 1 of 2.80 ^∘^C and at Fp1 electrode of 0.80 ^∘^C.

**Figure 8:**
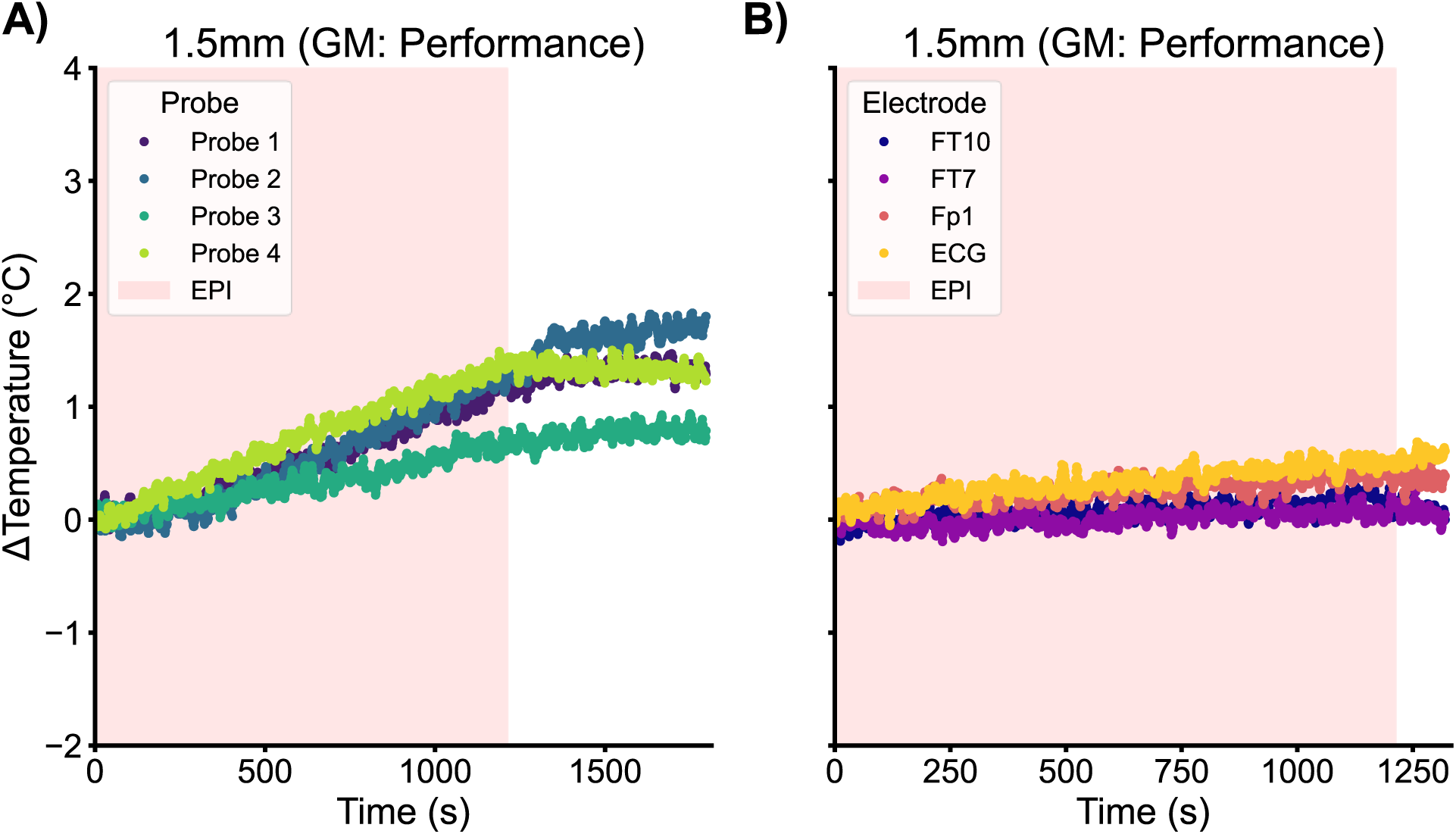
3T Temperature recordings from the amplifiers and powerpacks (panel **A**) and electrodes (panel **B**) when the High-Resolution (1.5 mm) EPI sequence was run for 20 mins with the 30 cm connector cable and the amplifiers in the ‘Sub-Optimal’ configuration. Pink shaded area represents EPI acquisition period. GM = gradient mode. Colours denote location of the temperature probes for panel **A**, as defined in the legend and Figures 2E & 2F.

## Discussion

The MRI scanner is a challenging place to record EEG data safely. Here, we demonstrate that when amplifiers and powerpacks are close to isocentre then the switching of gradients is sufficient to cause heating of amplifiers at 7T and 3T with different MR manufacturers. We show that at 7T the product EPI sequence has a high enough slew rate to cause heating of the EEG equipment when positioned directly behind the head coil. We then show that heating is reduced with reduced slew rates, through customisation of the MR sequence.

Furthermore, we showed that at 3T during EPI acquisition heating of the EEG amplifiers was considerable if the amplifiers were placed behind the head coil using short (10 cm) connector cables. With a High-Resolution EPI sequence over a 75 s recording, there was ∼3^∘^C temperature increase to the surface of an amplifier regardless of the configuration of the amplifiers and powerpacks (i.e. ‘Optimal’ vs ‘Sub-Optimal’). We estimated that this would result in the amplifiers heating by ∼62 ^∘^C over a 40 min run of the High-Resolution EPI sequence (Table S5), comparable to the duration of a standard EEG-fMRI study. This heating was reduced by moving the amplifiers further from the isocentre along the z-direction. With a 15 cm increase in distance from the head coil (achieved by making all cables taut), the predicted maximum heating over a 40 min period was reduced to ∼21 ^∘^C (Table S6). With a 37 cm increase (achieved by using 30 cm connector cables and making all cables taut), no measurable heating was observed (Table S7 C S8).

The change in heating with both sequence parameters and amplifier position provides evidence the heating is due to the gradient switching rather than to RF heating. The reduced number of slices in the High-Resolution sequence (22 vs 57) means that the SAR is lower than the Normal and Standard EPI sequences, because of the reduced number of excitation pulses and lower MB factor. A lower SAR would point to reduced chance of RF heating from the 3T High-Resolution sequence employed here. However, the High-Resolution sequence generated the strongest gradient flux due to its longer echo train length (High-Resolution: 123, Standard: 84, Normal: 84), readout time (High-Resolution: 94.52 ms, Standard: 48.96 ms, Normal: 48.96 ms), and the use of the performance gradient mode (maximum slew rate of 188.68 (T/m/s)). Thus, the greater heating observed for the High-Resolution compared with Normal MR sequences at 3T is consistent with the heating being driven by gradient switching rather than increased B1 fields. Further at 7T a birdcage transmit coil was used, thus the EEG amplifiers were well outside the transmit RF field, while the amplifier position is in a region of high magnetic field change from the gradient switching, providing further reason to suggest the heating observed is generated by the gradients.

When considering only the High-Resolution sequence, the heating reduced as the amplifiers were moved further from the isocentre in the z-direction. This may appear to be surprising; however, the maximum change in magnetic field with time (dB/dt) will be experienced ∼25-40 cm from isocentre, based on standard gradient performance with maximum amplitude per axis (40–80 mT), fastest increase time (100–400 μs), and maximum slew rate (130–200 T/m/s) (Stafford, 2020). We estimate that when the amplifiers are positioned behind the head coil, they are ∼34 cm from isocentre. This distance is then increased to ∼49 cm with the taut 10 cm connector cables and ∼71 cm with the taut 30 cm connector cables. Since the maximum dB/dt falls off rapidly with z-position outside the 25-40 cm “maximum exposure” area, this is likely to explain the changes in heating with position that we observed. Together our findings support that the heating observed in the amplifiers during EPI is driven by the gradient switching.

It is important to note that all our heating measurements were taken from the outside casing of the hardware. It is likely that whilst some of the heating will have occurred close to the plastic casing in the Faraday cage around the amplifier, the heating is greater inside the amplifier box where there is no air flow. However, it is also likely that the components of the amplifier are not heating to the level of the Faraday cage surrounding the amplifier within the plastic casing. With our measurements, it is not possible to know the precise temperatures that were reached within the amplifiers. However, we suggest that the amplifier and powerpack heating observed when they were positioned close to the back of the head coil present significant risks regardless of sequence and amplifier configuration used (of those tested here). Equipment damage may occur if the amplifier and/or powerpack reach temperatures above 50 ^∘^C (Brain Products GmbH, 2025). In addition, heating over an EEG-fMRI session may result in melting of the outer plastic case. These different safety concerns lead us to suggest that it is important to position the amplifiers as far from iso-centre as possible, whilst not neglecting the risks of RF antenna heating.

We therefore advise that it is very important to consider the experimental setup in the MRI scanner. The recommendation to use shorter connector cables by Brain Products was aimed at reducing the chance of RF burns to the participant (Brain Products GmbH, 2025), particularly taking into account the increase in SAR of sequences (e.g. multiband) now routinely used for EEG-fMRI. This safety consideration should not be under-estimated. However, when using the shorter connector cables, where the amplifiers are positioned relative to the largest changes in dB/dt from the gradient switching must also be carefully considered during experimental set-up. We recommend that heating tests be conducted at both the electrodes and amplifiers for each precise experimental setup, as gradient performance and B_1_ power will vary across MRI vendors and sequences.

We have estimated heating increases over continuous 40-minute EPI runs, which are unlikely to be used experimentally. However, 40 minutes of EPI data or more is commonly acquired during an EEG-fMRI session, with different paradigms or multiple runs of a single paradigm, e.g. (Griffiths et al., 2019; Mayhew, Hylands-White, et al., 2013; Mayhew, Ostwald, et al., 2013; Mullinger et al., 2013; Uji et al., 2018), or in relation to sleep (Hale et al., 2016; Jacob et al., 2025; Uji et al., 2025) and epilepsy (Coan et al., 2016; Flanagan et al., 2014) recordings. Since Figures 4-6 show that after the EPI sequences end the cooling of the amplifiers is very slow, it is likely that the heating effects would accumulate to something close to the temperature increases that are estimated for continuous 40-minute scans in this study.

One limitation of our analysis is that the majority of the scans carried out were of short duration; and from this we have estimated the heating over 40 minutes. During each acquisition, the RF and gradient conditions, equipment position, cable configuration, and airflow were held constant. Under these conditions, the electromagnetic power deposition was not expected to increase substantially with temperature, whereas heat dissipation would normally increase as the temperature difference between the equipment and the surrounding air increased. Therefore, the linear extrapolation is as a conservative upper-bound estimate of the temperature rise under the tested conditions and heating may have been somewhat less (at the points of measurement). However, given the sharp rise in heating observed, the inability to measure inside the amplifiers and the desire not to damage equipment 40-minute scans were not carried out for safety reasons.

In conclusion, we have demonstrated that amplifier and powerpack heating can be a direct result of the gradient fields, and that the heating rate is decreased by decreasing the gradient read-out slew rates or increasing the distance of the amplifiers from the maximum dB/dt. Using 30 cm connector cables reduced the heating on the EEG equipment by an order of magnitude, allowing longer EPI recordings without risk of damaging the equipment. Additionally, reducing the read-out gradient slew-rate, if the pulse sequence allows, will also substantially reduce the risk of damaging the EEG equipment (at the cost of a longer TR, but comparable image quality (Waggoner et al, 2024)). With these cables for the sequences we tested there was no evidence of RF heating on the EEG electrodes. Overall, we recommend careful consideration of the z-direction distance between the amplifiers and MRI scanner’s isocentre and slew rate, to avoid the risk of both equipment damage and to ensure participant safety.

## Ethics

All experiments were performed on phantoms. No human participants or animals were involved, and ethical approval was therefore not required.

## Acknowledgments

This work was supported by funding from BBSRC grant (BB/X013634/1), JSPS KAKENHI Grant Number JP22H01107 (MT), JP22K18664 (MT), JP25H00584 (MT), JP25K21987 (MT), JP26H00442 (MT), Naito Foundation (MT), Uehara Memorial Foundation (MT), and Takeda Science Foundation (MT).

## Data and Code Availability

All code and data can be found on GitHub: (https://github.com/Multimodal-Integration-Group/Ingram2026_EEGfMRIGradientHeating/).

The modified EPI sequence used for the Signa 7T data can be made available by the RIKEN authors to sites that possess a GE Healthcare Research Community Sharing License (RCSL).

3T Sequence parameters can be found in the Supplementary Information.

