## Supplementary Information for "Reducing the Risk of Gradient Induced EEG Hardware Heating During Simultaneous EEG-fMRI"

#### SAR Metrics

***Table S1:*** *GE Console reported SAR estimates for the 4 sequences used for 7T tests.*

| Sequence | *B1+RMS (uT)* | *SAR Head (W/Kg)* | *SAR Head (%)* | *Specific Energy Dose (J/KG)* |
|---|---|---|---|---|
| **EPI Slew 120** | 1.45 | 0.078 | 2.4 | < 0.1 |
| **EPI Slew 88** | 1.37 | 0.070 | 2.1 | < 0.1 |
| **EPI Slew 61** | 1.28 | 0.062 | 1.9 | < 0.1 |
| **EPI Slew 46** | 1.21 | 0.054 | 1.7 | < 0.1 |

**Table S2:** Siemens Console reported SAR estimates for the 3 sequences used for 3T tests.

| Sequence | B1+RMS (uT) | SAR Whole Body (W/Kg) | SAR Exposed Body (W/Kg) | SAR Head (W/Kg) | SAR HEAD local (%) | SAR Torso Local (%) | SAR Legs Local (%) | Specific Energy Dose (J/KG) | Time Averaged RF Power (W) |
|---|---|---|---|---|---|---|---|---|---|
| **High-Resolution** | 0.6 | 0 | 0.1 | 0.1 | 2 | 0.3 | 0.3 | 5 | 0.5 |
| **Standard** | 0.8 | 0 | 0.1 | 0.1 | 4.3 | 0.7 | 0.7 | 7 | 1 |
| **Normal** | 0.8 | 0 | 0.1 | 0.1 | 4.3 | 0.7 | 0.7 | 8 | 1 |

#### 7T Heating Results

***Table S3:*** *Temperature metrics for the 7T acquisitions. The Temperature probe position at the top of the EEG amplifier is defined in Figure 2B.*

| Sequence | Probe | Max ΔT (°C) | Slope (°C/min) | Extrapolated ΔT at 40 min (°C) |
|---|---|---|---|---|
| EPI Slew 120 | Probe 1 | 2.12 | 0.64 | 24.5 |
| EPI Slew 88 | Probe 1 | 1.46 | 0.41 | 17.5 |
| EPI Slew 61 | Probe 1 | 0.24 | 0.16 | 5.9 |
| EPI Slew 46 | Probe 1 | No heating | No heating | No heating |

### 3T Heating Results

***Table S4:*** *3T Temperature metrics per probe for each EPI sequence with 10 cm connector cables and the amplifiers and powerpacks in the 'Optimal' configuration. Temperature probe positions are defined in Figures 2C&D.*

| Sequence | Probe | Max ΔT (°C) | Slope (°C/min) | Extrapolated ΔT at 40 min (°C) |
|---|---|---|---|---|
| High-Resolution | Probe 1 | 2.71 | 1.76 | 70.40 |
| | Probe 2 | 2.12 | 0.84 | 33.60 |
| | Probe 3 | 0.02 | No Heating | No Heating |
| | Probe 4 | 2.94 | 1.64 | 65.60 |
| Standard | Probe 1 | 1.61 | 0.95 | 38.00 |
| | Probe 2 | 1.07 | 0.50 | 20.00 |
| | Probe 3 | 0.06 | No Heating | No Heating |
| | Probe 4 | 1.70 | 0.98 | 39.20 |
| Normal | Probe 1 | 1.46 | 0.95 | 38.00 |
| | Probe 2 | 0.98 | 0.35 | 14.00 |
| | Probe 3 | 0.06 | No Heating | No Heating |
| | Probe 4 | 1.53 | 1.09 | 43.60 |

***Table S5:*** *3T Temperature metrics per probe for each EPI sequence with 10 cm connector cables and the amplifiers and powerpacks in the 'Sub-Optimal' configuration. Temperature probe positions defined in Figures 2E&F.*

| Sequence | Probe | Max ΔT (°C) | Slope (°C/min) | Extrapolated ΔT at 40 min (°C) |
|---|---|---|---|---|
| High-Resolution | Probe 1 | 2.19 | 1.30 | 52.00 |
| | Probe 2 | 0.11 | No Heating | No Heating |
| | Probe 3 | 0.75 | 0.25 | 10.00 |
| | Probe 4 | 3.04 | 1.56 | 62.40 |
| Standard | Probe 1 | 1.51 | 0.88 | 35.20 |
| | Probe 2 | 0.26 | No Heating | No Heating |
| | Probe 3 | 0.60 | 0.06 | 2.40 |
| | Probe 4 | 1.59 | 1.04 | 41.60 |
| Normal | Probe 1 | 1.35 | 1.20 | 48.00 |
| | Probe 2 | 0.31 | No Heating | No Heating |
| | Probe 3 | 0.54 | 0.06 | 2.40 |
| | Probe 4 | 1.28 | 0.76 | 30.40 |

***Table S6:*** *3T Temperature metrics per probe for each EPI sequence with 10 cm connector cables pulled taut and the amplifiers and powerpacks in the 'Sub-Optimal' configuration. Temperature probe positions defined in Figures 2E & 2F.*

| Sequence | Probe | Max ΔT (°C) | Slope (°C/min) | Extrapolated ΔT at 40 min (°C) |
|---|---|---|---|---|
| High-Resolution | Probe 1 | 1.03 | 0.53 | 21.20 |
| | Probe 2 | 0.12 | No Heating | No Heating |
| | Probe 3 | 0.27 | No Heating | No Heating |
| | Probe 4 | 1.08 | 0.28 | 11.20 |
| Standard | Probe 1 | 0.52 | 0.16 | 6.40 |
| | Probe 2 | 0.03 | No Heating | No Heating |
| | Probe 3 | 0.22 | No Heating | No Heating |
| | Probe 4 | 0.52 | 0.12 | 4.80 |
| Normal | Probe 1 | 0.32 | No Heating | No Heating |
| | Probe 2 | 0.15 | No Heating | No Heating |
| | Probe 3 | 0.45 | No Heating | No Heating |
| | Probe 4 | 0.61 | 0.10 | 4.00 |

***Table S7****: 3T Temperature metrics per probe for each EPI sequence with 30 cm connector cables and the amplifiers and powerpacks in the 'Sub-Optimal' configuration. Temperature probe positions defined in Figures 2E & 2F.*

| Sequence | Probe | Max ΔT (°C) | Slope (°C/min) | Extrapolated ΔT at 40 min (°C) |
|---|---|---|---|---|
| High-Resolution | Probe 1 | 0.13 | No Heating | No Heating |
| | Probe 2 | 0.05 | No Heating | No Heating |
| | Probe 3 | 0.07 | No Heating | No Heating |
| | Probe 4 | 0.29 | No Heating | No Heating |
| Standard | Probe 1 | 0.16 | No Heating | No Heating |
| | Probe 2 | 0.05 | No Heating | No Heating |
| | Probe 3 | 0.12 | No Heating | No Heating |
| | Probe 4 | 0.18 | No Heating | No Heating |

***Table S8:*** *3T Temperature metrics per probe for the High-Resolution sequence with 30 cm connector cables and the amplifiers and powerpacks in the 'Sub-Optimal' configuration for a 20-minute recording. Temperature measurements were acquired on the amplifiers and powerpacks, as well as the EEG electrodes during a separate EPI acquisition. Temperature probe positions on amplifier and powerpacks are defined in Figures 2E & 2F.*

| Sequence | Probe | Max ΔT (°C) | Slope (°C/min) | Extrapolated ΔT at 40 min (°C) |
|---|---|---|---|---|
| High-Resolution (Amplifiers & powerpacks) | Probe 1 | 1.43 | 0.06 | 2.40 |
| | Probe 2 | 1.78 | 0.07 | 2.80 |
| | Probe 3 | 0.89 | 0.03 | 1.20 |
| | Probe 4 | 1.46 | 0.07 | 2.80 |
| High-Resolution (EEG electrodes) | FT10 | 0.25 | 0.01 | 0.40 |
| | ECG | 0.16 | 0.01 | 0.40 |
| | FT7 | 0.44 | 0.01 | 0.40 |
| | Fp1 | 0.64 | 0.02 | 0.80 |
